# 3’ untranslated regions of Marburg and Ebola virus mRNAs possess negative regulators of translation that are modulated by ADAR1 editing

**DOI:** 10.1101/2021.04.21.440871

**Authors:** Sudip Khadka, Caroline G. Williams, Joyce Sweeney-Gibbons, Christopher F. Basler

## Abstract

The filovirus family includes deadly pathogens such as Ebola virus (EBOV) and Marburg virus (MARV). A substantial portion of filovirus genomes encode 5’ and 3’ untranslated regions (UTRs) of viral mRNAs. Select viral genomic RNA sequences corresponding to 3’UTRs are prone to editing by ADAR1. A reporter mRNA approach, in which different 5’ or 3’UTRs were inserted into luciferase encoding mRNAs, demonstrates that MARV 3’UTRs yield different levels of reporter gene expression suggesting modulation of translation. The modulation occurs in cells unable to produce miRNAs and can be recapitulated in a minigenome assay. Deletion mutants identified negative regulatory regions at end of the MARV NP and L 3’UTRs. Apparent ADAR1 editing mutants were previously identified within the MARV NP 3’UTR. Introduction of these changes into the MARV nucleoprotein (NP) 3’UTR or deletion of the region targeted for editing enhances translation, as indicated by reporter assays and polysome analysis. In addition, the parental NP 3’UTR, but not the edited or deletion mutant NP 3’UTRs, induce a type I interferon (IFN) response upon transfection into cells. Because some EBOV isolates from the West Africa outbreak exhibited ADAR1 editing of the VP40 3’UTR, VP40 3’UTRs with parental and edited sequences were similarly assayed. The EBOV VP40 3’UTR edits also enhanced translation but neither the wildtype nor the edited 3’UTRs induced IFN. These findings implicate filoviral mRNA 3’UTRs as negative regulators of translation that can be inactivated by innate immune responses that induce ADAR1.

**Importance:** UTRs comprise a large percentage of filovirus genomes and are apparent targets of editing by ADAR1, an enzyme with pro- and antiviral activities. However, the functional significance of the UTRs and of ADAR1 editing have been uncertain. This study demonstrates that MARV and EBOV 3’UTRs can modulate translation, in some cases negatively. ADAR1 editing or deletion of select regions within the translation suppressing 3’UTRs, relieves the negative effects of the UTRs. These data indicate that filovirus 3’UTRs contain translation regulatory elements that are modulated by activation of ADAR1, suggesting a complex interplay between filovirus gene expression and innate immunity.

## Introduction

The family *Filoviridae*, which includes Ebola virus (EBOV) and Marburg virus (MARV), is comprised of filamentous, enveloped, negative-sense RNA viruses (1). Several filovirus family members are zoonotic pathogens that cause sporadic outbreaks associated with high case fatality rates and efficient human-to-human transmission (2). Examples include the West Africa epidemic from 2014-2016 that was associated with more than 28,000 cases and 11,000 deaths (3), an EBOV outbreak in the Democratic Republic of Congo from 2018-2020 with more than 3,400 cases and nearly 3,000 deaths, and a 2005 MARV outbreak in Angola with a reported 88% case fatality rate (4).

The approximately 19 kilobase long MARV and EBOV genomic RNAs serve as templates for viral genome replication, which yields viral anti-genomic and genomic RNAs, and for transcription, where seven genes serve as separate transcription units that produce mRNAs encoding viral proteins. MARV and EBOV mRNAs are 5’-capped, 3’ polyadenylated and possess long untranslated regions (UTRs) (5). The UTRs are predicted to possess secondary structures (6, 7). The MARV Angola strain genome (KU978782.1) encodes 5’UTRs ranging from 54-108 nucleotides and 3’UTRs ranging from 302-684 nucleotides in length. Overall, these comprise 22.2% of the genome. UTRs occupy a similar percentage of the EBOV genome. This contrasts with vesicular stomatitis virus (VSV), a representative negative-sense RNA virus of the rhabdovirus family, which has 5’ and 3’UTRs that account for 1.0% and 2.5% of the genome, respectively.

The functions of filovirus UTRs are incompletely understood but include impacts on viral transcription and mRNA translation. In one well-defined example, a stem-loop near the EBOV nucleoprotein (NP) mRNA transcriptional start site makes EBOV transcription dependent on the viral VP30 protein (5). The impact of different EBOV 5’UTRs on mRNA translation has also been surveyed through the use of *in vitro* transcribed and transfected model mRNAs (8). This implicated short upstream open reading frames (uORFs) present in the 5’UTRs of the EBOV VP35, VP30, VP24 and L mRNAs as cis-acting regulators of protein synthesis. The L uORF was shown to modulate L mRNA translation such that, when cell stress was low, the uORF substantially attenuated L translation; however, when stress and eIF-2α phosphorylation increased, L translation was upregulated (8).

Sequences within the negative-sense RNA genomes that correspond to 3’UTRs appear to be targets of editing by adenosine deaminases acting on RNA 1 (ADAR1), enzymes that catalyze the deamination of adenosine (A) to inosine (I) in dsRNA structures (9). Such editing was suggested by RNAseq studies of MARV-Angola infected cells with U→C mutations accumulating in select 3’UTRs of viral mRNAs, most dramatically in the MARV NP 3’UTR (10). This suggests deamination of adenosine (A) to inosine (I), with I being the functional equivalent of guanosine (G), leading to the U→C changes in the positive-sense mRNA. In a separate study, serial passage in mice of the MARV-Angola strain led to 26 A→G changes that accumulated in the negative-sense genome in regions encoding the NP mRNA 3’UTR (11). In another example, mouse-adaptation of Ravn virus, which represents a distinct clade within the genus *Marburgvirus*, led to the accumulation of 30 A→G changes within sequences corresponding to the 600 nucleotide long 3’UTR of the glycoprotein (GP) mRNA (12). During the 2014-2016 West Africa EBOV outbreak varied A→G changes were identified in the negative-sense viral RNAs of different isolates, with sequences encoding the VP40 3’UTR being a hotspot for such changes (13-19). What functional impact ADAR1 editing of 3’UTRs may have on EBOV and MARV replication is unclear.

The present study addresses the functional significance of filovirus 3’UTRs and the editing of these sequences by ADAR1. The data demonstrate that the UTRs of MARV Angola regulate translation by miRNA-independent mechanisms. A region that suppresses translation is identified within the MARV NP 3’UTR and this corresponds to a region of the viral genome previously implicated as a target for editing by ADAR1. ADAR1 editing mutations or deletions of this region enhance expression from model mRNAs. Whereas the unedited MARV NP 3’UTR activates the IFNβ promoter, the edited or deleted 3’UTR does not. The enhanced expression is recapitulated using a minigenome assay and reflects enhanced translation as demonstrated by polysome assays. Finally, the roles of 3’UTRs and ADAR1 editing are extended to EBOV by studying parental and edited 3’UTRs corresponding to the VP40 gene of West Africa isolates. These data provide functional insight into filoviral UTRs and suggest novel mechanisms regulating filoviral gene expression.

## Results

### MARV 3’ UTRs regulate mRNAs

The relative lengths of MARV and EBOV 5’UTRs, coding sequences and 3’UTRs are indicated in Fig. 1A. In order to investigate the role of MARV UTRs on mRNA translation we generated reporter constructs with an individual MARV 3’UTR and/or 5’UTR flanking the *Renilla* luciferase coding sequence. These were placed adjacent to the T7 promoter sequence (Fig 1B). The constructs were PCR amplified and used for in vitro transcription. Each resulting RNA was capped and polyA tailed. Equal amounts of RNAs were then transfected into HEK293T cells and luciferase activity was measured at early (2h) and late (20h) times post transfection. The constructs with the NP and VP35 5’UTRs showed modestly higher luciferase signals at both the early and later time points, as compared to the other constructs (Fig 1C), suggesting an enhancing role of these 5’UTRs. The remaining mRNAs with MARV 5’ UTRs yielded very similar levels of expression. With the 3’UTR containing constructs, there were varying degrees of expression. The NP, GP, VP24 and L 3’UTR containing constructs exhibited lower levels of expression as compared to the others, whereas the VP30 3’UTR containing construct gave the highest levels of expression at an earlier time point (Fig 1D). We next used reporter-mRNAs with both the 5’ and 3’ UTRs from each viral gene to better mimic the MARV mRNAs. The VP35, VP30 and VP24 signals showed higher levels of luciferase activity at earlier time points (Fig. 1E) whereas the NP, VP40, GP and L respectively exhibited a decreasing order of activity. The lower levels of expression observed with the GP and L UTR containing mRNAs did not recover at later time points. Overall, these results suggest that the 5’ and 3’ UTRs of MARV mRNAs regulate translation and, because the expression patterns for the mRNAs with only 3’UTRs parallel those with both 5’ and 3’UTRs, the 3’ UTRs exert dominant effects.

**Figure 1:**
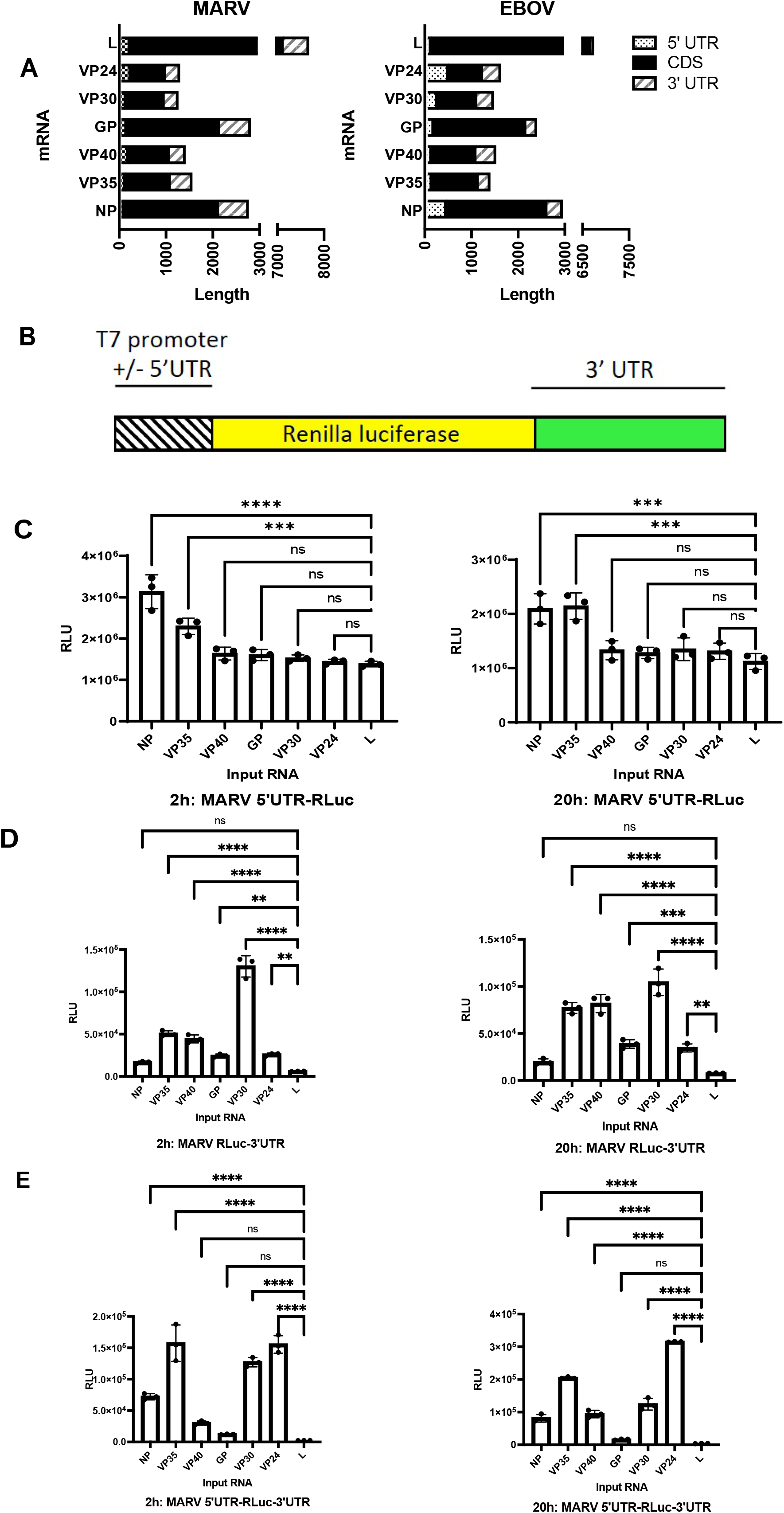
Filovirus mRNA UTRs modulate translation efficiency. (A) Depiction of the relative length of MARV and EBOV mRNA 5’UTRs, coding sequences and 3’UTRs. NP, nucleoprotein; VP35, viral protein of 35 kDa; VP40; GP, glycoprotein; VP30, VP24 and L; large protein mRNAs. (B) Test mRNAs were constructed by flanking the Renilla luciferase coding sequences with the MARV 5’ and/or 3’UTRs. (C-E) Renilla luciferase activities 2- or 20-hours post-transfection of test mRNAs. Test mRNAs with 5’UTR alone (C), 3’UTR alone (D) and both the 5’ and 3’ UTRs (E) were evaluated. The data represent the mean and standard deviation (SD) of triplicate samples (*P<0.05, **P<0.01, ***P<0.001, ****P<0.0001).

### MARV 3’UTRs regulate expression independently of miRNAs

MicroRNAs (miRNAs) regulate gene expression through mechanisms that include modulating mRNA stability and mRNA translation, often through targeting of mRNA 3’UTRs (20). To determine whether miRNAs mediate the changes in reporter gene expression mediated by the MARV UTRs, we compared expression from the reporter mRNAs in HEK293T cells or HEK293T-derived RNAse III knockout cells which lack both Drosha and Dicer and therefore are defective for miRNA production (21, 22). Both cell types were transfected side-by-side with the 5’UTR-RLuc (Fig. 2A), RLuc-3’UTR (Fig. 2B) or 5’UTR-RLuc-3’UTR (Fig. 2C) mRNAs. Luciferase activity was measured at 24 hours post transfection to allow for sufficient time for any miRNA mediated action on the transfected mRNAs. In each instance, reporter gene expression in RNAse III knock out cells mirrored that in HEK293T cells, suggesting that the miRNAs do not play a significant role in translational regulation of the MARV mRNAs. Upon comparison of normalized RLU values between the HEK293T cells and the RNAse III KO cells, the only significant differences were modest and found in the mRNA with NP 5’UTR (Fig. 2A) or both the 5’ and 3’ UTR containing VP24 and VP35 test mRNAs (Fig 2C). None of the test mRNAs containing only the 3’UTR showed any significant changes in expression levels between the two cell lines, suggesting that the miRNAs play minimal, if any, role in 3’UTR mediated translational regulation of MARV genes.

**Figure 2:**
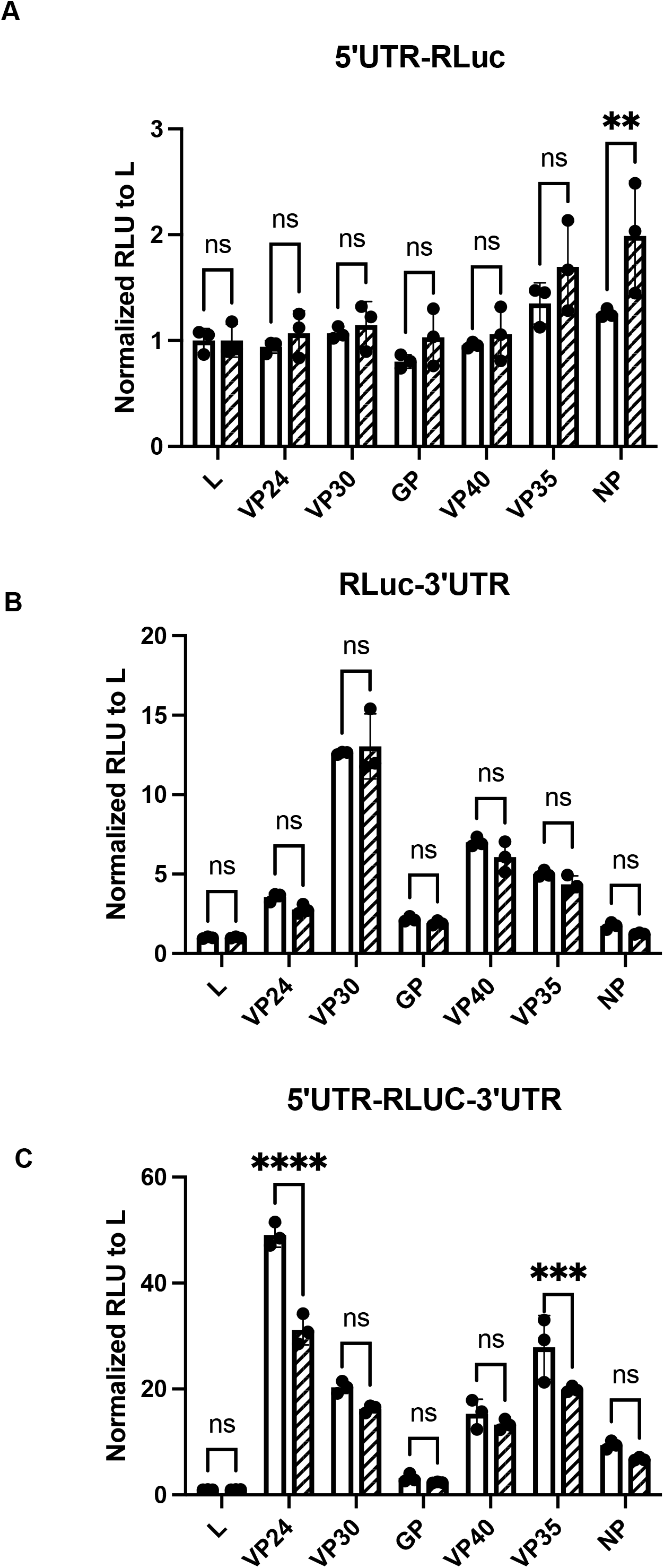
Modulation of translation by filoviral UTRs is not dependent on the action of miRNAs. HEK293T-derived RNAse III knockout cells that are defective in miRNA production were transfected with test mRNAs and luciferase levels were measured 4 hours post transfection. Test mRNAs had MARV (A) 5’UTRs, (B) 3’UTRs and (C) both 5’ and 3’UTRs. RLU values were normalized to the L UTR containing mRNA expression values in each set. The data represent the mean and SD of triplicate samples (*P<0.05, **P<0.01, ***P<0.001, ****P<0.0001).

### The MARV NP 3’UTR contains negative regulatory elements

MARV mRNAs possess a conserved sequence of AUUAAGAAAA, corresponding to the viral gene end signal, at their 3’ ends of the UTRs (Fig. 3A). Preserving this conserved sequence, we deleted approximately 100 bases from the 3’ end for each of the RLuc-3’UTR mRNAs (Fig. 3B). We then assessed the expression of *Renilla* luciferase 2h post transfection. A substantial increase in expression from the mutated NP, VP30 and L truncation constructs was seen. The L truncation construct (L1-420) exhibited an 18-fold increase in luciferase signal (Fig. 3B). The length of a 3’UTR can modulate mRNA translation (23). To test whether the enhancement of translation seen in our truncation mutants was specifically due to length of the 3’UTR, we generated a series of additional NP 3’UTR mutants with successive truncations of roughly one hundred nucleotides (Fig 3C). *In vitro* transcribed mRNA from these constructs were then transfected into the HEK293T cells and analyzed for expression of luciferase (Fig. 3D). Reduction of 3’UTR length from 654 to 523 nucleotides resulted in enhanced expression. However, larger truncations did not further increase expression until the 3’UTR was reduced to 78 nucleotides, where another bump in expression was noted. We also analyzed the expression of the constructs at later time points and saw no change in the pattern of expression (Fig. 3D). These data demonstrate that luciferase expression levels are not strictly dependent upon the length of the 3’UTR and suggest that the 3’ end of the NP 3’UTRs contains a negative regulatory element(s).

**Figure 3:**
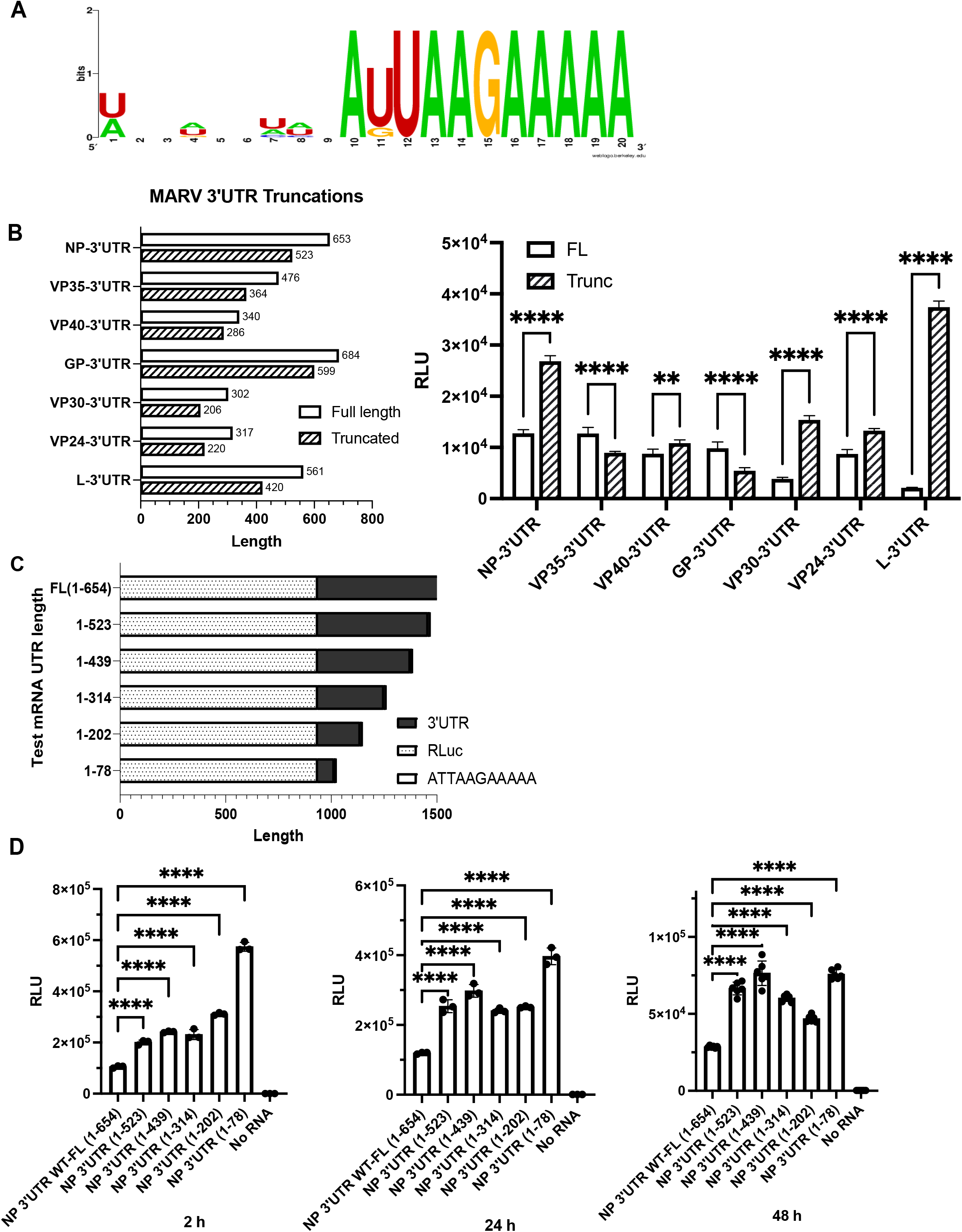
MARV NP 3’UTR contains negative regulatory sequences. (A) Alignment of the last 20 bases of 3’UTRs illustrate the conserved transcription stop signal AU/GUAAGAAAAA found at the ends of all the MARV 3’UTRs. (B) Left. A roughly 100 base deletion was introduced to each of the MARV 3’UTRs, as illustrated, while preserving the conserved sequence at the end. Right. Luciferase activities following transfection of the full length and truncated 3’UTR test mRNAs. (C) Illustration of further truncations made to the NP 3’UTR. (D) Luciferase activities after 2, 24 or 48 hours following transfection of test mRNAs with the 3’UTRs illustrated in panel C. The data represent the mean and SD of triplicate samples except for the 48h samples where sextuplicate samples were assayed (*P<0.05, **P<0.01, ***P<0.001, ****P<0.0001).

### MARV NP 3’UTR mutations due to apparent ADAR1 editing of the MARV genome enhance mRNA translation efficiency

A previous study in which mRNAs from MARV-infected THP1 and Vero cells were analyzed by RNAseq at 12 and 24 hours post infection identified ten mutations in the NP 3’UTR (10). These all clustered in the second half of the NP 3’ UTR (Fig. 4A). To study the effect of the putative ADAR1 editing mutations, we generated reporter constructs with either the wild type or the 3’UTR with the complete set of editing mutations. Each condition was tested in the context of an mRNA that possessed or lacked the NP 5’ UTR. Equal amounts of *in vitro* transcribed RNAs were transfected into HEK293T cells and luciferase activity was measured 6h later. Expression from the mRNAs with the mutated 3’UTR were higher than the wild type 3’UTR mRNAs, regardless whether the NP 5’UTR was present (Fig. 4B). However, there was increased expression from the 5’UTR containing mRNAs as compared to those lacking the 5’UTR. The enhancing effect of the 3’UTR mutations was evident as early as 4 hours and as late as 48 hours post transfection (Fig. 4B).

**Figure 4:**
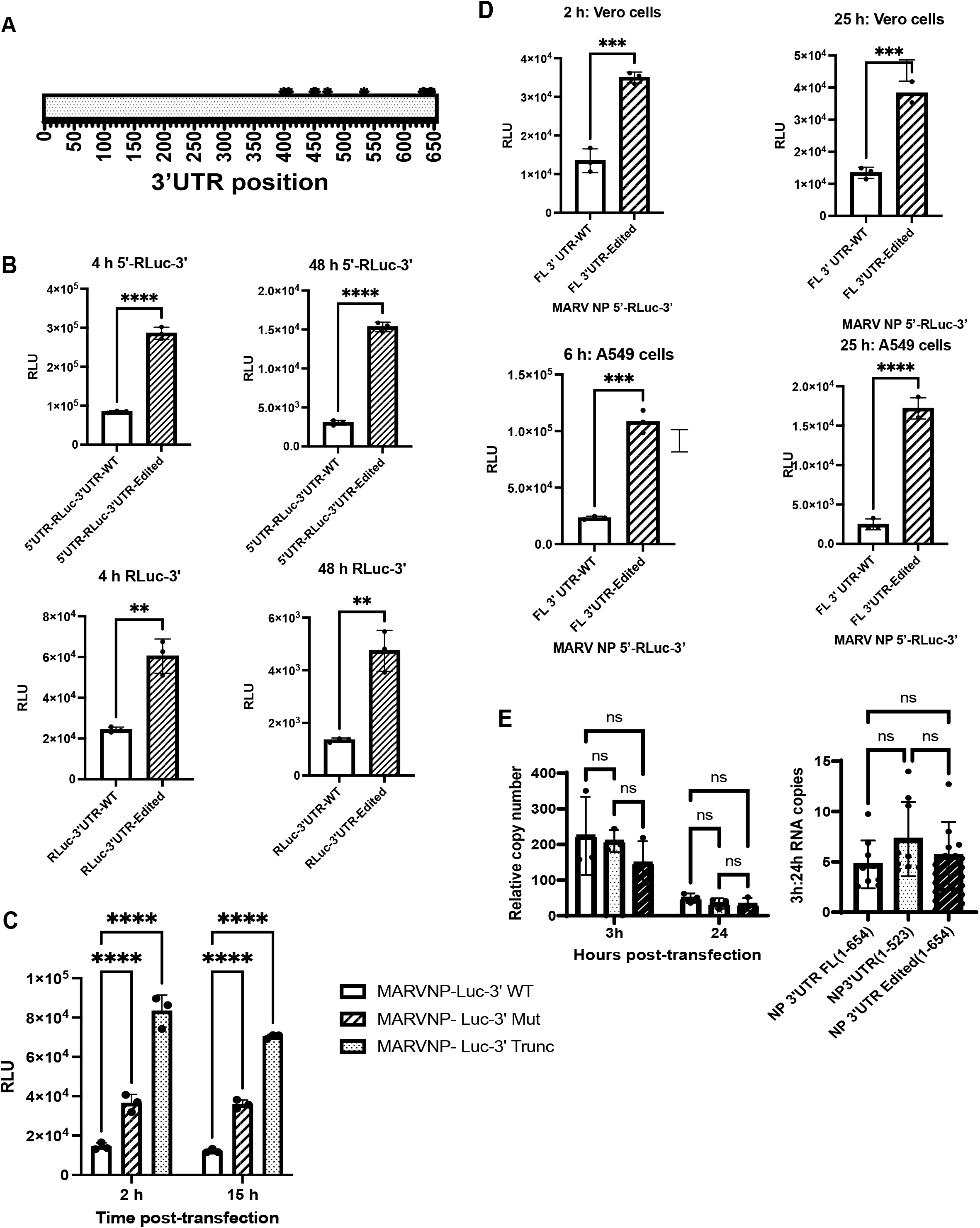
Edits observed in the MARV NP 3’UTR relieve translational suppression. (A) Schematic of the MARV NP 3’UTR showing positions of the editing mutations. All the changes map to the second half of the NP 3’UTR with 9 changes from U to C and one change from A to G. (B) Renilla luciferase expression at 4 hours and 48 hours post transfection of test mRNAs with wildtype (WT) or edited 3’UTRs. The test mRNAs possessed (top graphs) or lacked (lower graphs) the 5’UTRs. (C) *Renilla* luciferase activities at 2 and 15 hours post-transfection of Vero cells with WT, edited or 3’ truncated 3’UTR (1-395) test mRNAs. (D) Comparison of test mRNAs with WT and edited 3’UTRs in Vero cells (top graphs) versus A549 cells (bottom graphs) at 2 and 25 hours post-transfection. The data represent the mean and SD of triplicate samples (*P<0.05, **P<0.01, ***P<0.001, ****P<0.0001). (E) Relative copy numbers at 3 and 24 hours post-transfection (left) and the ratio of 3 hour to 24 hours mRNA copy numbers (right) following transfection of wt, truncated and 3’UTR edited test mRNAs. The amount of transfected RNA present in the cells was determined by reverse transcription-qPCR and normalized to human GAPDH mRNA levels. Rate of decay of the WT, truncated or the edited mRNAs were not statistically different (ns) from one another.

Additional constructs were made where the MARV NP 3’UTR region with the observed editing mutations was deleted. This region encompassing the region 1-395 of the 3’UTR was used to study the effect of deletion of the edited region. Transfection of the Vero cells showed that the mRNAs with only the first 395 bases of the 3’UTR had a significant increase in translation at both early and later time points (Fig. 4C). This bump in translation observed was even greater than that observed for the editing mutations. We also analyzed enhancement of translation by each of the ten individual mutations. However, we did not observe the same levels of enhancement due to any single mutation suggesting that more than one change is required to see the enhancement in translation (data not shown).

Comparison of these mRNA constructs in Vero cells, which do not produce type I interferons (IFN) and A549 cells which do, yielded similar results, at both early and late time points, demonstrating that the enhancing effect of the mutations is not unique to a given cell type and suggesting that it is not directly related to the IFN response (Fig. 4D).

Stability of mRNAs can dictate the level of translation. To determine whether increased translation from the mutated RNAs was due to altered RNA stability, we measured the ratio of RNA levels at early (3h) and late (24h) timepoints post-transfection by qRT-PCR (Fig. 4E). Ratios of copy numbers at 3 hours and 24 hours post transfection were found to be similar for all the transfected mRNAs suggesting a very similar rate of mRNA decay. We also included a truncated 3’UTR (1-523) that showed a significant increase in translation compared to the full-length 3’UTR (Fig. 3D) as an additional control. No significant differences in the ratio of early to late mRNAs were detected, suggesting that the higher expression from the mutated or truncated NP 3’UTRs is not due to increased mRNA stability.

### Enhancement of translation due to edits in the 3’UTR is independent of the identity of the 5’UTR

Interactions between 5’UTR and 3’UTR sequences, due either to direct RNA-RNA pairing or via RNA binding proteins, can influence translation efficiency. For our model MARV mRNAs, the presence of the NP 5’UTR did not alter the enhancement due to the 3’UTR mutations (Fig. 4B). To further address possible impacts of the 5’UTRs, we replaced the MARV 5’UTR with 5’UTRs from three different human genes ACE, ABHD11 and ACO2 in the context of mRNAs with either the NP WT or edited 3’UTRs (Fig 5A). These 5’UTRs were chosen to represent a shorter (ACE, 22 bases), same length (ABHD11, 50 bases) or longer (ACO2, 321 bases) 5’UTR, respectively (Fig. 5A). Following transfection into Vero cells, the mRNAs with different 5’UTRs yielded different levels of luciferase expression, suggesting that each individual 5’UTR regulates translation to different extents. However, in each case, irrespective of the 5’UTR present, the mutated 3’UTR yielded higher luciferase activity (Fig. 5B). These results reinforce the view that mutations in the 3’UTR enhance translation in a 5’UTR independent manner.

**Figure 5:**
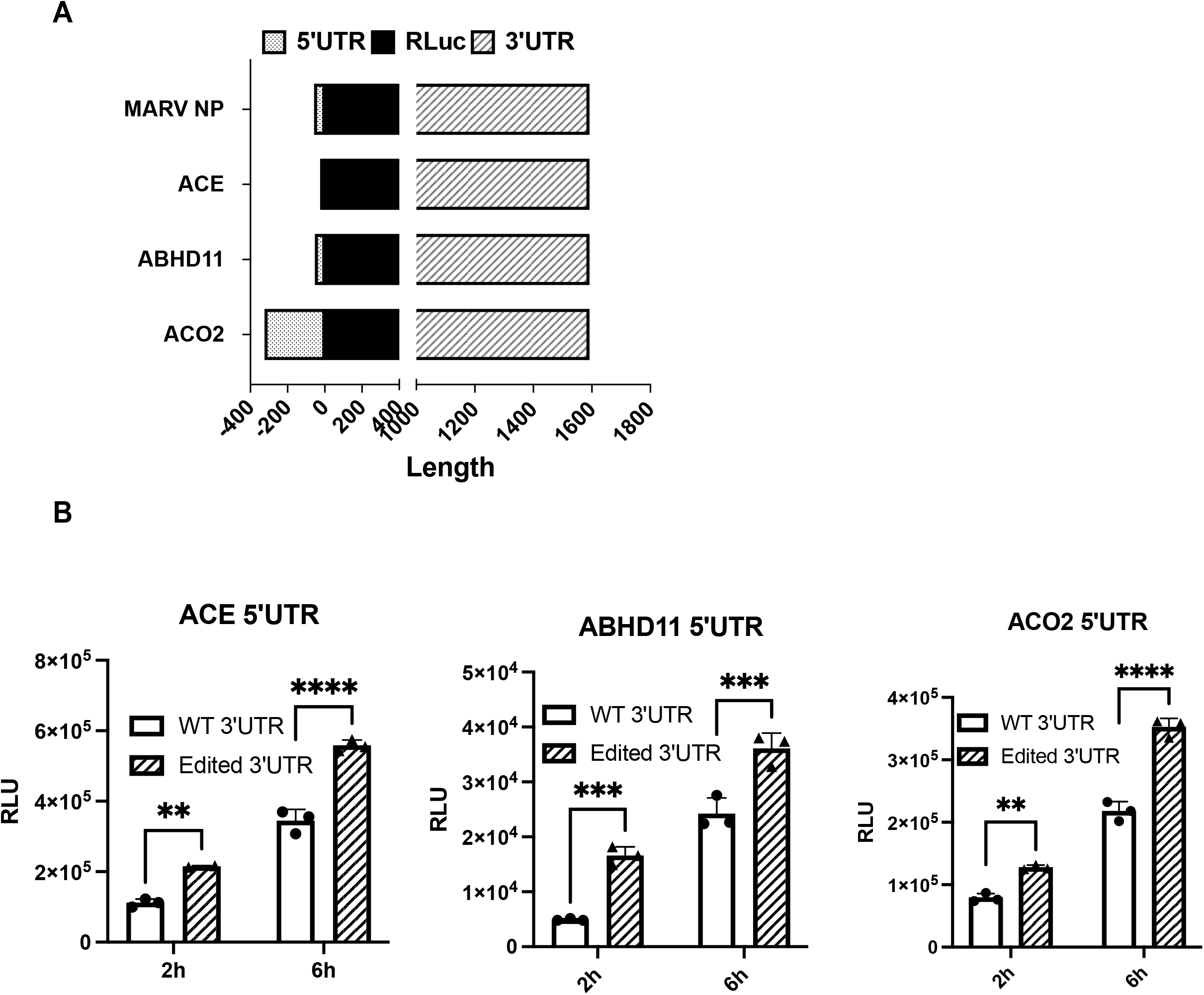
Enhancement of translation due to edits in the NP 3’UTR are independent of the 5’UTR. (A) Illustration of mRNAs possessing 5’UTRs from MARV NP, ACE, ABHD11 and ACO2 and the WT and edited MARV NP 3’UTR and encoding *Renilla* luciferase (RLuc). (B) *Renilla* luciferase activities 2h and 6h post transfection of Vero cells with the test mRNAs illustrated in (A). The data represent the mean and SD of triplicate samples (*P<0.05, **P<0.01, ***P<0.001, ****P<0.0001).

### Mutations in the MARV NP 3’UTR of the MARV NP mRNA relieves an interferon-inducing activity

The recognition of the MARV 3’UTR by ADAR1 suggests that it might also be recognized by other pattern recognition receptors. To determine whether mRNAs with NP 5’ or 3’UTRs can induce a type I IFN response, we utilized a stable cell line that has firefly luciferase under control of the human IFNβ promoter. These cells were transfected with capped, polyadenylated and phosphatase treated *Renilla* luciferase reporter mRNAs. Activation of the IFNβ promoter was assessed by measuring the firefly luciferase activity, whereas *Renilla* luciferase activity gave a measure of translation from the input RNA. Transfected were mRNAs that either did or did not possess the NP 5’UTR and had the wildtype NP 3’UTR, a 3’UTR with the ADAR1 editing mutations, no 3’UTR or a truncated 3’UTR. Whereas the mRNAs with the wildtype 3’UTR activated the IFNβ promoter, the remaining mRNAs elicited a substantially reduced response (Fig. 6A). The wildtype 3’UTR construct induced the most IFN response and was translated to the lowest levels (Fig. 6B). We carried out similar experiments utilizing a stable cell line that expresses firefly luciferase under control of a promoter with an IFN-stimulated response element (ISRE) and obtained similar results (Fig. 6C). We also measured by quantitative RT-PCR levels of endogenous mRNAs, including those for IFN-β and IFN stimulated genes (ISGs) upon transfection with these RNAs (Fig. 6C). The IFN-β, RIG-I, TNF-α, ISG 15, IRF1, MxA, STAT1 and ISG56 mRNA levels were found to be higher in cells transfected with the WT 3’UTR as compared to the mutant 3’UTR or an mRNA lacking UTRs. Levels for TBK1 mRNA, although higher for WT UTR transfected cells, were not found to be significantly different between the two conditions tested. These data indicate that the NP 3’UTR contains negative regulatory elements that reduce gene expression but can activate innate antiviral defenses. The ADAR1 mutations relieve the repression, resulting in enhanced translation of the gene product while reducing innate immune response.

**Figure 6:**
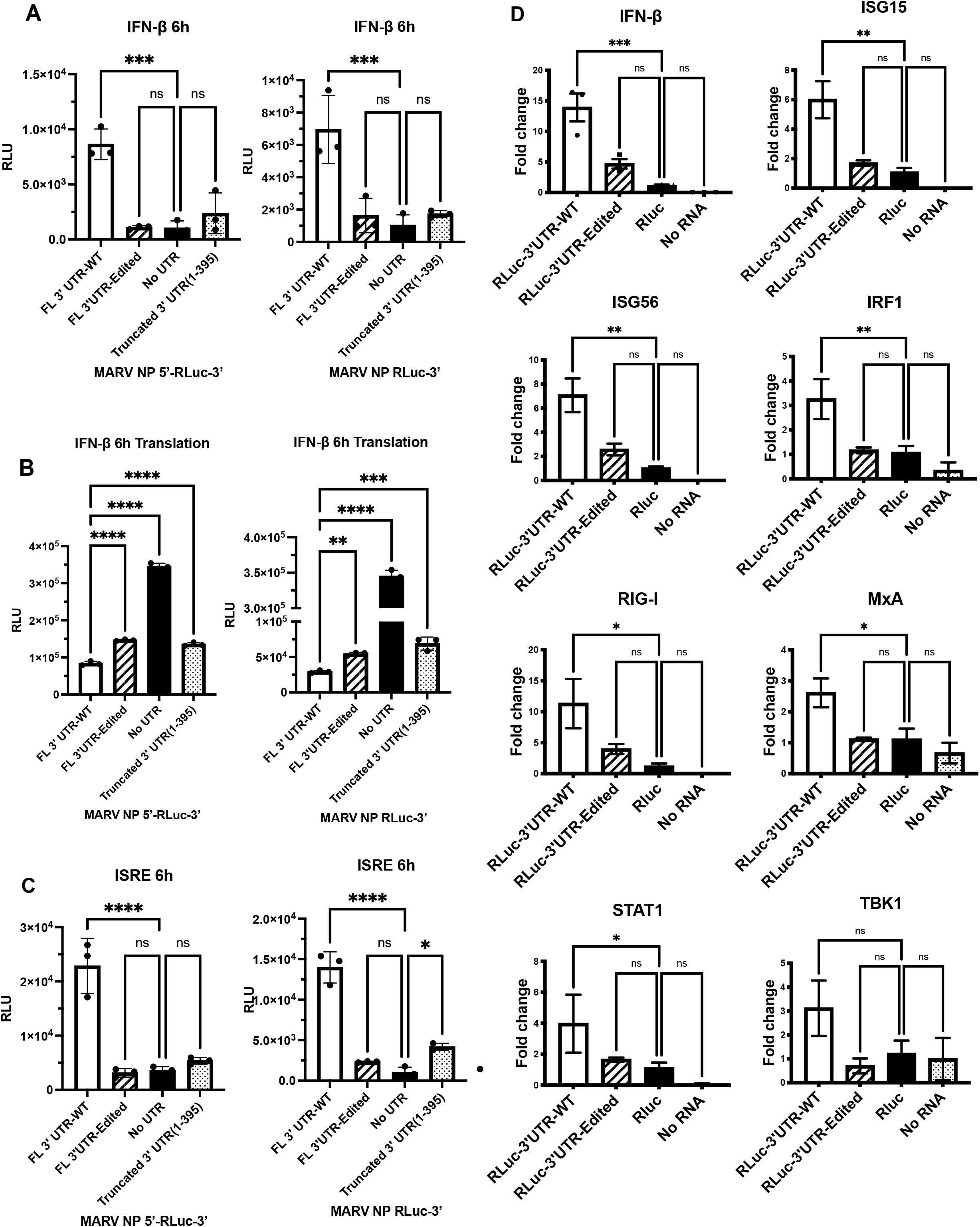
Edits in the MARV NP 3’UTR relieve an IFN inducing activity. (A) Firefly luciferase activity expressed from a stable IFN-β promoter-luciferase cell line following transfection of test mRNAs with the WT, edited or truncated (1-395) NP 3’UTRs at 6h post transfection. (B) Renilla luciferase activity, produced from the transfected test RNAs described in panel A. (C) Stable cell lines with firefly luciferase reporter under control of an Interferon Stimulated Response Element (ISRE) promoter were similarly transfected as in panel A and assayed for activation at 6h post transfection. (D) IFN and IFN stimulated gene (ISG) expression following transfection of A549 cells with the indicated test mRNAs. IFN-β, ISG15, ISG56, IFN regulatory factor 1 (IRF1), retinoic acid inducible gene I (RIG-I), MX dynamin like GTPase (MxA), signal transducer and activator of transcription 1 (STAT1), and Tank binding kinase 1 (TBK1) mRNAs were measured qRT-PCR. Levels were normalized to GAPDH mRNA and Renilla luciferase (RLuc) mRNA without UTRs as control. All test RNAs were capped, polyadenylated and phosphatase treated before transfection. The data represent the mean and SD of triplicate samples (*P<0.05, **P<0.01, ***P<0.001, ****P<0.0001).

### Incorporation of equivalent mutations in the genomic strand of a model viral RNA results in higher levels of protein expression

Since it appears that ADAR1 editing occurs on the MARV negative sense genomic RNA, we constructed a bicistronic minigenome system that incorporates the NP 3’UTR. The bicistronic minigenome was designed to code for model NP and L mRNAs where the coding sequences were replaced with *Renilla* and firefly luciferase coding sequences, respectively (Fig. 7A). Negative strand minigenome RNA was *in vitro* transcribed, purified and then transfected into the cells 24 hours after transfection of cell with helper plasmids encoding MARV NP, VP35, VP30 and L, which transcribe and replicate the minigenome RNA. A further 24 hours after transfection of the minigenome RNA, *Renilla* luciferase activity was measured to assess the impact of changes in the NP 3’UTR. Similar to what was observed with the mRNA transfections, the minigenome encoding a mRNA that has mutations in the NP 3’UTR had higher luciferase activity compared to the one encoding the WT NP 3’UTR (Fig. 7B). The controls lacking the VP35 or the L helper plasmids did not show any luciferase activity, demonstrating that active replication and transcription from transfected minigenome RNA was necessary for expression of the reporter gene.

**Figure 7:**
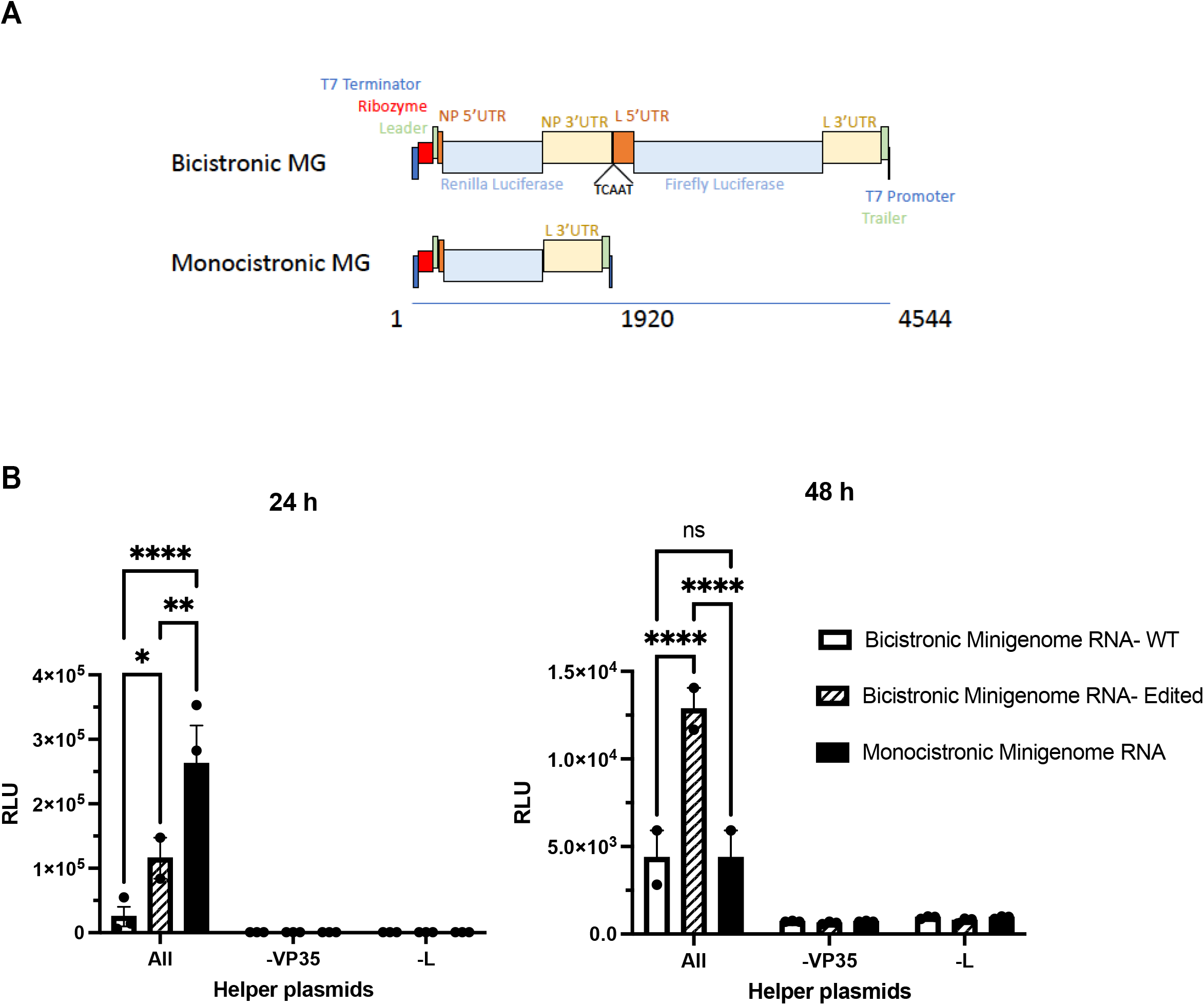
Incorporation of NP 3’UTR editing mutations increases expression from a bicistronic minigenome. (A) A bicistronic minigenome (MG) (top) was designed such that the *Renilla* luciferase (RLuc) coding sequence was flanked by the MARV NP 5’UTR and either the wt or edited NP 3’UTR. (Bottom) illustration of a monocistronic MG encoding RLuc. (B) Renilla luciferase activities at 24 or 48 hours post transfection of the indicated bicistronic or monocistronic minigenome RNAs. Helper plasmids were transfected 24 hours prior to transfecting the MG RNAs. The data represent the mean and SD of triplicate samples (*P<0.05, **P<0.01, ***P<0.001, ****P<0.0001).

### mRNAs with mutations in the 3’UTR associate better with the polysomes

To further assess how mutations in the NP 3’UTR modulate translation, polysome analysis was performed. HEK293T cells were transfected with purified mRNA possessing WT or mutant NP 3’UTRs. The cells were harvested 4 h post transfection after a brief treatment with cycloheximide to immobilize the ribosomes on actively translating mRNAs. The cleared lysate was then separated on a continuous sucrose density gradient by ultracentrifugation and fractions were collected. Absorbance profiles (260 nm) of the flow during fractionation were taken to identify the polysome containing fractions. RNA was then extracted from all the fractions and analyzed by qRT-PCR to quantify the amount of transfected RNA present. Compared to the mRNA with WT 3’UTR mRNA (Fig. A), a larger proportion of the mRNA with mutated 3’UTR associated with the polysome fractions (Fig. 8B). Increased association of the edited NP 3’UTR containing mRNA with polysomes is in agreement with our previous observation that the 3’UTR mutations enhance translation efficiency.

**Figure 8:**
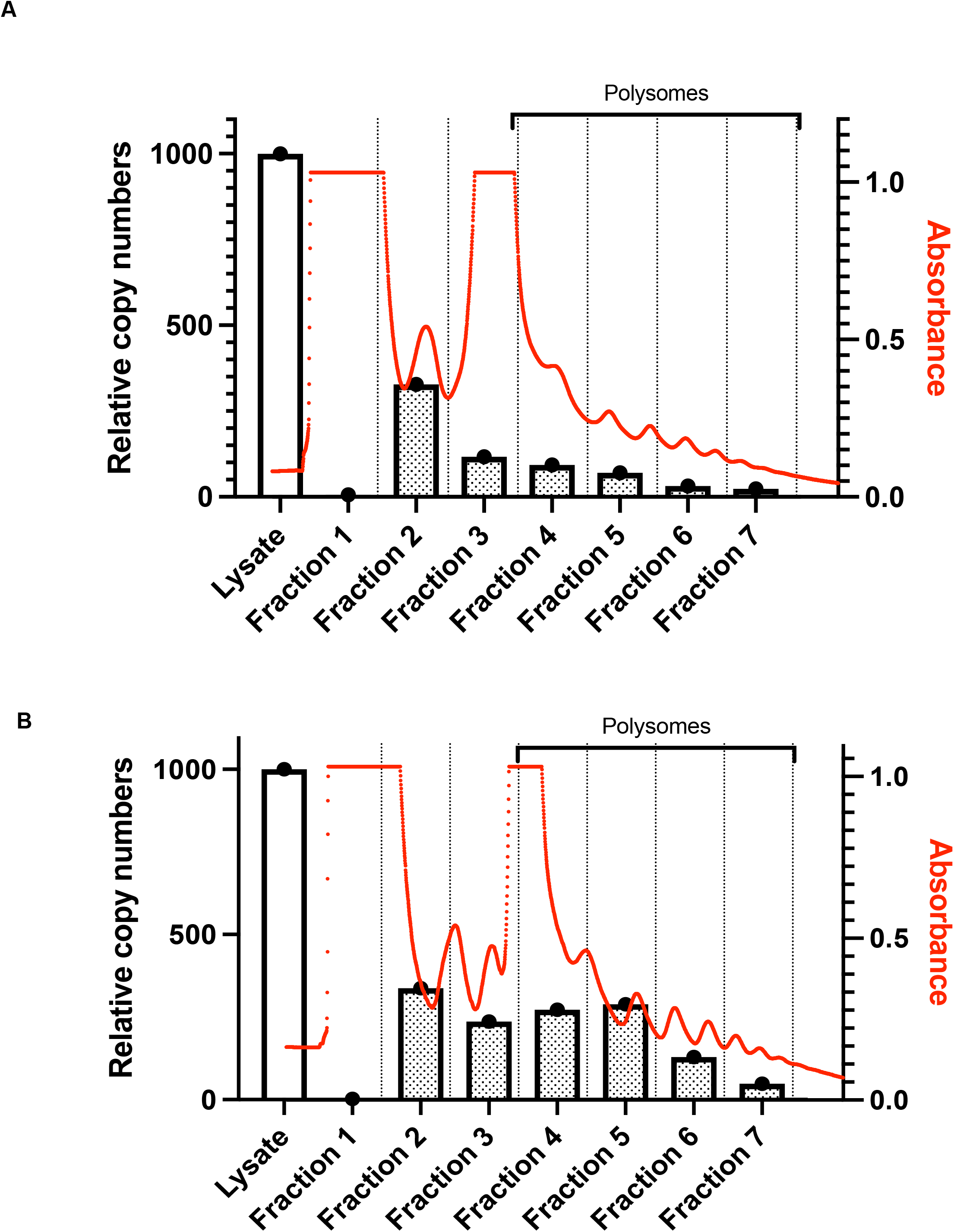
mRNAs with edits in the NP 3’UTR are enriched in polysome fractions compared to the WT 3’UTR containing mRNAs. Relative copy numbers of test mRNAs present in cell lysates and in fractions following polysome analysis (left side y-axis) and absorbance at 260 nm (right side y-axis). Following transfection of the HEK293T cells with test mRNAs, ribosomes were immobilized on the actively translating mRNAs by treating with cycloheximide. Cells were lysed, separated on a sucrose density gradient and fractioned. Total RNA was extracted from each fraction and analyzed by reverse transcription-qPCR using primers for RLuc to determine distribution of transfected test mRNA in each fraction. Comparison of (A) WT NP 3’UTR containing test mRNAs and (B) edited 3’ UTR containing test mRNA.

### Apparent ADAR editing of the VP40 3’UTR from the Ebola virus isolated from the 2014-2016 outbreak results in increased translation

During the 2014-2016 West Africa EBOV outbreak varied A→G changes were identified in the negative-sense viral RNAs of different isolates, with sequences encoding the VP40 3’UTR being a hotspot for such changes (13-19). A comprehensive analysis of 1086 publicly available full-length EBOV-Makona genome sequences identified 49 with clusters of A→G substitutions (24). Of these 30 had A→G clusters in sequences corresponding to 3’UTRs, and fifteen of these had clusters in sequences corresponding to the VP40 3’UTR. Within this group, eleven isolates had 12-13 A→G changes (24). Analysis of intra-patient virus sequences documented conversion of 13 A→G changes during the course of disease in two different patients, demonstrating that these mutations can arise independently in different individuals (24). In total, 15 different ADAR1 mediated mutations were described, with fourteen of these from T to C and one change from G to A (Fig. 9A). We incorporated these changes into the EBOV VP40 3’UTR and compared expression to that of the WT VP40 3’UTR sequence using our reporter system. Similar to what was observed for the MARV NP 3’UTR, the edited EBOV VP40 3’UTRs yielded higher translation efficiency at both early (2h) and later (18h) time points (Fig. 9B). To examine RNA stability, the ratio of RNA in the cells at earlier and later timepoints was calculated. The ratios were similar for the WT and edited 3’UTR containing constructs, suggesting a very similar decay rate for both the RNA species (Fig. 9C). This suggests that the increased translation is not attributable to the stability of the RNAs.

**Figure 9:**
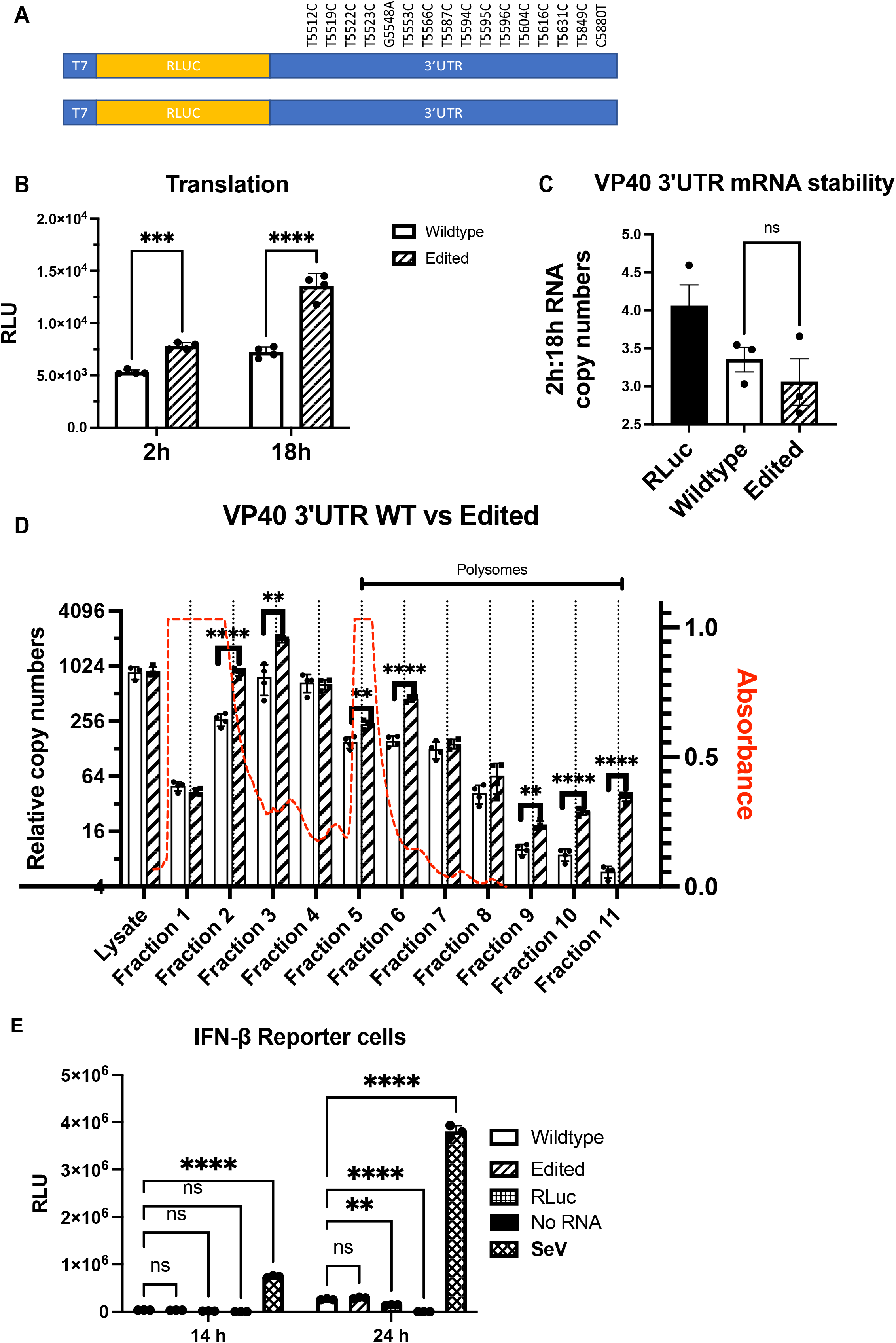
Edits identified in the 3’UTR of EBOV Makona VP40 mRNA enhance translation. (A) Design of test mRNAs with EBOV VP40 3’UTR with editing mutations (top) or WT sequence (bottom). The positions of edits in the EBOV genome are indicated. RLuc, *Renilla* luciferase. (B) *Renilla* luciferase activities 2 and 18 hours post-transfection of the WT and edited test mRNAs. (C) The ratio of WT and edited EBOV VP40 3’UTR test mRNA copy numbers at 2h and 18h post transfection. The amounts of transfected mRNA present in cells. Copy number was determined by normalizing the test mRNA levels to GAPDH mRNA levels. (D) Polysome analysis of WT and edited EBOV VP40 3’UTR test mRNAs. Wildtype test mRNA levels are depicted by white bars and edited test mRNAs by hatched bars. The analysis was performed as described in Fig. 8. (E) Firefly luciferase activities following transfection of WT and edited EBOV VP40 3’UTR test mRNAs into the stable IFN-β promoter-firefly luciferase reporter cells. The experiment was performed as described for Fig. 6A and B. The data represent the mean and SD of triplicate samples except for (B) where quadruplicate samples were used (*P<0.05, **P<0.01, ***P<0.001, ****P<0.0001).

To directly assess effects of 3’UTR mutations on translation, polysome analysis was performed. A higher abundance of the edited 3’UTR mRNA in the polysome fractions as compared to the WT 3’UTR mRNA (Fig 9D). This suggests that like the MARV 3’UTR, the EBOV VP40 mRNA 3’UTR contains elements that impair translation and that these can be relieved by ADAR1 editing. We also assessed activation of the IFN-β reporter gene with these mRNAs in reporter cells. Unlike the MARV NP 3’UTRs, we did not observe differential induction of the IFN-β promoter by the edited mRNAs (Fig 9E). The positive control Sendai virus infection robustly activated the promoter at both 14 and 24 hours post infection.

## Discussion

Given the minimal information available regarding the functional role of filovirus UTRs, this study sought to determine how filovirus UTRs modulate the translation of mRNAs. In order to make comparisons between different MARV UTRs, the model mRNAs encoding *Renilla* luciferase were flanked by MARV 5’UTRs, 3’ UTRs or both 5’ and 3’UTRs. After *in vitro* transcription, capping and polyadenylation, the mRNAs were purified and transfected into cells. Using luciferase activity as a measure of gene expression at different time points post-transfection, we characterized the impact of both the 5’ and 3’ UTRs on expression. Comparing the translation efficiency of the model MARV mRNAs with both 5’ and 3’UTRs identified differences in expression for each model mRNA. Expression increased in the following order: L, GP, VP40, NP, VP30, VP35 and VP24. When we compared the reporter mRNAs containing only the 5’UTR we observed that the expression levels for all mRNAs were comparable although the NP and the VP35 5’UTR mRNAs yielded slightly higher levels than the rest. This suggested that the variation in translation efficiency in the MARV mRNAs was mostly due to the 3’UTRs. When constructs with only the MARV 3’UTRs were compared, varying effects on translation were obtained. The NP, GP, VP24 and L 3’ UTR containing mRNAs showed lower expression compared to the rest. Interestingly, the L 3’UTR yielded the least signal, suggesting limited expression of the L protein in infected cells. Low expression of protein translated from mRNAs with MARV L UTRs parallels the observation that EBOV L mRNA is maintained at low levels by cis-acting elements present in the L mRNA 5’UTR (8). Notably, translation efficiency did not correspond to the lengths of either the 5’ or 3’UTRs. To further refine where regulatory elements reside, model mRNAs with 3’UTRs truncated from the 3’ end were produced. Significant enhancement was demonstrated following transfection of truncated NP and L 3’UTRs. To further address the possibility that the 5’UTRs impact the effects of the 3’UTRs, swaps of MARV 5’UTRs with unrelated human 5’UTRs of varying lengths were made. These changes did not substantially alter the modulating effects of the 3’UTRs.

UTRs can act as determinants of mRNA stability with various sequences imparting longer or shorter half-life for the mRNAs. Short AU rich elements (AREs) present in 3’UTRs are well documented to regulate mRNA stability in the eukaryotic cells (25). Structural elements present within the 3’UTRs also play a role in mRNA stability. For example, the constitutive decay element present in the tumor necrosis factor 3 (TNF3) mRNA forms a stem-loop structure that is recognized by Roquin and Roquin2 proteins which then recruit Ccr4-Caf1-Not deadenylase complex for initiation of mRNA decay (26). The Zc3h12a endonuclease recognizes a stem-loop structure in the 3’UTR of Interleukin (IL)-6 and regulates its decay (27). To address possible impacts of the 3’UTRs on the stability of the transfected mRNAs, we harvested cells at early and later time points, determined the amount of each transfected mRNA present in the cells by qRT-PCR and the ratio of transfected mRNA in at early and late time points was then calculated to determine the rate of decay. Comparable rates of decay were observed for the mRNAs tested, suggesting they have very similar stabilities, regardless of whether the 3’UTR sequence was wild type or edited. Therefore, mRNA stability is unlikely to explain differences observed in translation.

A major mechanism regulating mRNA translation in eukaryotic cells is by the action of miRNAs. While coding sequences and both the 5’ and 3’ UTRs in host and viral mRNAs are known to harbor miRNA target sites, it is the 3’UTRs where such target sites are most abundant (28-30). The complex relationship between the host miRNAs and the viral RNA is well documented in several viruses, where negative and positive effects have been demonstrated (31-34). To address the possible role of miRNAs in the effects of the MARV 3’UTRs, transfections were performed in cells possessing an intact miRNA biogenesis system and in a RNAse III knockout cell line that lacks both Dicer and Drosha rendering them defective in miRNA biogenesis (21). In these cells, no difference in the translation regulation by either the 5’ or the 3’ UTR was detected, ruling out an essential role for miRNAs in regulation of translation mediated by the UTRs in our assays. The negative regulatory elements in the NP 3’UTR correspond to sites in the MARV genome that appear to be edited by ADAR1 (10). ADAR1 enzymes catalyze the deamination of adenosine (A) to inosine (I) in dsRNA structures (9). A-to-I RNA editing in the context of cytoplasmic RNA viruses has been attributed to ADAR1, which has two isoforms, p110 and p150. p110 is constitutively expressed and mostly nuclear, whereas p150 is IFN-inducible and mainly cytoplasmic. ADAR1 prevents inappropriate activation of the innate immune system by editing cytoplasmic RNAs, primarily pol II-transcribed Alu elements, that otherwise activate MDA5 (35-40). ADAR1 also suppresses activation of the IFN-induced, dsRNA-activated antiviral kinase, PKR; edits RNA viruses and can act in a pro-viral manner (41-43).

ADAR1 editing was suggested by RNAseq studies of MARV-Angola infected cells with sites of U→C mutations accumulating in select 3’UTRs of viral mRNAs, most dramatically in the MARV NP 3’UTR. In THP-1 cells, at 24 hours post-infection, 7 sites with U→C changes in 9-35% of reads were identified (10). This suggests deamination on the negative-sense viral genomic RNA of adenosine (A) to inosine (I), with I being the functional equivalent of guanosine (G), leading to the U→C changes in the positive-sense mRNA. When these mutations were built into model mRNAs, the impairment of reporter gene expression was relieved, mirroring the effect of deletions in the 3’UTR. It is notable that similar editing also occurred during serial passage in mice of the MARV-Angola strain, where 26 A→G changes accumulated in the negative-sense genome in regions encoding the NP mRNA 3’UTR (11). These are not the only sites in MARVs that appear to be susceptible to such editing. Mouse-adaptation of Ravn virus, which represents a distinct clade within the genus *Marburgvirus*, led to the accumulation of 30 A→G changes within sequences corresponding to the 600 nucleotide long 3’UTR of the GP mRNA (12). It will be of interest to determine if these editing events also impact translation of the targeted mRNAs and whether this facilitates adaptation to the mouse.

ADAR1 also appears to act on EBOV. During the 2014-2016 West Africa EBOV outbreak varied A→G changes were identified in the negative-sense viral RNAs of different isolates, with sequences encoding the VP40 3’UTR being a hotspot for such changes (13-19). During the 2014-2016 West Africa EBOV outbreak varied A→G changes were identified in the negative-sense viral RNAs of different isolates, with sequences encoding the VP40 3’UTR being a hotspot for such changes (13-19) (24). Of these 30 had A→G clusters in sequences corresponding to 3’UTRs, and fifteen of these had clusters in sequences corresponding to the VP40 3’UTR. Within this group, eleven isolates had 12-13 A→G changes (24). Analysis of intra-patient virus sequences documented conversion of 13 A→G changes during the course of disease in two different patients, demonstrating that these mutations can arise independently in different patients (24). Building these mutations into model mRNAs with EBOV VP40 3’UTRs yielded a similar outcome as with the MARV NP 3’UTR mRNAs, suggesting negative-regulatory elements in EBOV mRNAs as well. Given that ADAR1 targets dsRNA, these data suggest the presence of secondary structures that contribute to the negative effects of the 3’UTR on translation. These data also suggest the possibility that activation of innate antiviral responses which upregulate ADAR1 p150 may increase translation efficiency of some filovirus mRNAs. A caveat to these analyses is that not all sites are edited at the same efficiency. For edits identified in MARV infected Vero cells, at 24 hours post-infection, frequencies of U→C changes in the NP 3’UTR ranged from 8-24 percent. In infected THP-1 cells at the same time point, frequencies of mutations ranged from 9-35 percent (10). Further, the deep sequencing approaches that identified the ADAR1 edits do not determine which of the various changes are present on the same RNA molecule.

Editing of the viral genomic RNA by ADAR1 suggests the presence of secondary structures that other pattern recognition receptors might also recognize. We addressed this by asking whether wildtype and edited 3’UTRs, which would be complementary to the edited genomic RNA, might activate the interferon β (IFNβ) promoter or an IFN-inducible ISRE promoter. Our data demonstrate that 5’-capped, 3’-polyadenylated and phosphatase-treated mRNAs possessing the MARV WT NP 3’UTR but not the edited 3’UTR induced IFN responses, suggesting this UTR may contribute to IFN responses. Interestingly, a similar effect was not detected with the EBOV VP40 3’UTR. There is precedent for the activation of IFN responses by capped RNAs, as 3’UTRs from influenza virus and some RNA aptamers have been described as 5’-triphosphate-independent RIG-I activators (44, 45). There is also substantial evidence that ADAR1 editing serves as a mechanism to prevent induction of IFN responses and activation of PKR by self and viral RNAs (35, 36, 38, 40, 42, 46). Therefore, ADAR1 may be hypothesized to relieve the IFN response triggered by NP 3’UTRs during MARV infection. To what extent such an effect occurs in infected cells remains to be determined. The VP35 proteins of EBOV and MARV inhibit RIG-I mediated activation of IFN responses, so it is possible these activities would mask the effects of 3’UTR (47, 48).

The finding that filovirus 3’UTRs include elements that decrease translation efficiency is intriguing. That such elements can also activate innate immune responses, as in the case of the NP 3’UTR containing mRNAs, and be susceptible to modification by ADAR1, the cytoplasmic p150 form of which is IFN-inducible is also notable. These findings suggest that possible costs of these features are offset by other, yet to be identified, benefits. A previous study identified an upstream open reading frame in the EBOV L 5’ UTR as a suppressor of L translation, but the suppression could be relieved when phosphorylation of translation initiation factor eIF2a was induced. This was proposed as a mechanism by which L protein levels could be maintained at a low but consistent level, even when innate immune responses are activated (8). A similar phenomenon could be at play with the MARV NP 3’UTR or the EBOV VP40 3’UTR. Preferred levels of protein expression might be achieved by a less than maximum translation efficiency under some circumstances. Inclusion of a regulatory element that is inactivated when IFN responses are triggered and ADAR1 p150 expression is induced could ensure sustained expression of the viral mRNA despite activation of innate immunity. Such a mechanism might be most relevant early in infection when viral mRNA levels are lower. A key to evaluating these hypotheses will be testing these in infected cells.

## Materials and Methods

### Cell culture

A549 cells, human embryonic kidney (HEK)293T cells and Vero cells were obtained from ATCC. RNAse III knockout cells were a kind gift from Benjamin tenOever (Icahn School of Medicine at Mount Sinai, New York). Stable cell lines with IFNβ and ISRE reporters have been described previously (49). All cells were cultured in a temperature controlled humidified incubator maintained at 37°C and 5% CO_2_. Dulbecco’s Modified Eagle Medium (DMEM) with 10% fetal bovine serum (FBS) was used as growth medium.

### Synthesis of RNA constructs

Both wild type and edited MARV NP 3’UTR coding sequences were synthesized by Genscript (Piscataway, NJ). For amplifying the remaining MARV UTRs purified total RNA from MARV infected THP1 cells at 24 hours post infection was used for cDNA synthesis. The 5’ and 3’ UTR coding sequences were amplified with UTR-specific primers. Appearance of mutations in the 3’UTR coding region of Ebola virus VP40 gene during the course of infection were reported from patients in Sierra Leone during the 2014-2016 West Africa Ebola virus outbreak (24). A total of 16 different changes were observed in the VP40 3’UTR as compared to Zaire ebolavirus isolate H.sapiens-wt/GIN/2014/Makona-Kissidougou-C15 (GenBank accession No. KJ660346). Wildtype 3’UTR and 3’UTR coding sequences containing all 16 editing mutations were synthesized commercially (Genscript). Forward primers were designed to add the T7 promoter <u>gactcg</u>taatacgactcactataggggaagag at the 5’ end. *Renilla* luciferase (RLuc) coding sequence was flanked by either the 5’UTR, 3’UTR or both the UTRs to create templates for transcribing test mRNAs. Assembly of the RNA coding sequence and template was accomplished by PCR using Q5 High-Fidelity DNA polymerase (NEB). Finally, template DNA was amplified with the forward primer gactcgtaatacgactcactataggggaag and a 3’ end specific reverse primer. The amplified template was gel purified, *in vitro* transcribed using HiScribe™ T7 Quick High Yield RNA Synthesis Kit (NEB) using the manufacturer’s protocol, purified by LiCl_4_ precipitation and capped using the Vaccinia Capping System (NEB). Capped mRNAs were again purified by LiCl_4_ precipitation followed by the addition of polyA tails using E coli Poly(A) polymerase (NEB) as per the manufacturer’s protocol. After a final round of LiCl_4_ purification, mRNA was resuspended in nuclease free water, analyzed by formaldehyde agarose gel electrophoresis, aliquoted and stored at -20 °C. Truncated mRNAs were generated in a similar fashion by using the full length template for amplification. Reverse primers specific to internal sites were designed with a conserved terminal sequence ATTAAGAAAAA at the 3’end.

### mRNA reporter assay

*In vitro* transcribed RNA was transfected into cells using Lipofectamine 2000 reagent (Invitrogen). Briefly, cells were seeded in a T75 flask 24 hours prior to transfection such that they would be ∼80% confluent on the day of transfection. These cells were reverse transfected. The transfection used 10,000 cells and 100 ng of RNA per 96 well plate or scaled accordingly. At the specified times, levels of luciferase expression were measured using the Renilla-Glo™ Luciferase Assay System (Promega) per the manufacturer’s protocol.

### Quantifying Total RNA

Total RNA was prepared from transfected samples using Trizol (Invitrogen) reagent. cDNA synthesis was performed using oligo-dT primers for mRNA or random hexamers for total RNA using the SuperScript IV Reverse Transcriptase kit (ThermoFisher Scientific) as per the manufacturer’s protocol. Quantitative PCR (qPCR) was carried out following the manufacturer’s protocol using SYBR Green Master Mix (ThermoFisher) on a Biorad CFX real time PCR system (49). GAPDH mRNA, levels were quantified using specific primers (Fwd-CACCCACTCCTCCTACTTT, Rev-CCCTGTTGCTGTAGCCAAAT). Comparisons were made by the ddCT method. For rate of decay assays, total RNA was extracted at early (2h-4h) and late time points (18h-24h). Following cDNA synthesis using oligo dT primers, reporter mRNA was quantified was by using RLuc specific primers (Fwd AACGCGGCCTCTTCTTATTT, Rev ATTTGCCTGATTTGCCCATA). A ratio of RNA levels at early and later time points was taken as a measure of input RNA stability.

### Interferon β and ISRE promoter reporter assays

Stable cell lines with an IFNβ promoter reporter and an ISRE promoter reporter have been described previously (49). Both cell lines express firefly luciferase upon activation. mRNAs encoding *Renilla* luciferase were transfected into the stable reporter cells and were grown using standard methods. At the indicated time points firefly and *Renilla* luciferase activities were measured using the Dual-Glo® Luciferase assay system (Promega) following the manufacturer’s protocol. Student’s t-tests were carried out with GraphPad Prism to assess statistical significance.

### Minigenome assay

HEK 293T cells were grown in T75 flasks until they were ∼80% confluent on the day of transfection. Helper plasmids that express the MARV L, NP, VP30 and VP35 proteins have been previously described (50). These plasmids were transfected into the cells in T75 flasks. At 24 hours post transfection of helper plasmids, *in vitro* transcribed and purified minigenome RNA was reverse transfected into the cells, reseeded in 24 well or 96 well plates and the cells were grown for the indicated times. Minigenome assays was evaluated by measuring Renilla luciferase activity and by quantitative RT-PCR. Strand specific primers were used for cDNA synthesis from genomic sense (GGACACACAAAAAAGATGAAGAATG) or antisense (TGGACACACTAAAAAGATGAAGAATG) minigenome RNA strands. Following cDNA synthesis standard qPCR was done using the Renilla Luciferase primers.

### Bicistronic minigenome

A bicistronic minigenome was designed to contain two genes. The first gene possessed *Renilla* luciferase coding sequences flanked by the MARV NP 5’ and 3’UTRs and the second gene possessed firefly luciferase coding sequences flanked by the MARV L 5’ and 3’ UTRs. The region encoding wildtype or edited MARV NP 3’UTRs was PCR amplified from constructs used in the comparative expression assays. cDNA obtained from infected THP1 cells was used to amplify the MARV L 5’UTR. A ‘middle fragment’ was assembled by PCR and was comprised of the MARV NP 3’UTR- the intergenic sequence TCAAT- the L 5’UTR- and firefly luciferase. Similarly, a ‘front fragment’ comprised of T7 terminator-ribozyme-Leader-NP 5’UTR-RLuc and an ‘end fragment’ comprised of L 3’UTR-trailer-T7 promoter were PCR amplified from a monocistronic MARV minigenome template. Next, the front, middle and end fragments were assembled by overlapping PCR amplification. All PCRs were carried out using Q5 High-Fidelity polymerase (NEB). The resulting construct was designed to produce a genomic-sense RNA upon *in vitro* transcription. The PCR amplicons were gel purified, sequenced and used as a template for *in vitro* transcription. Transcribed bicistronic minigenome RNA was purified by LiCl_4_ precipitation and size verified by running on a formaldehyde agarose gel.

To confirm transcription of mRNAs containing the wild type and edited NP 3’UTRs in cells transfected with the bicistronic minigenome and helper plasmids, total RNA was harvested from transfected cells. cDNA was generated by using oligo-dT primers and quantitative PCR and sequencing was performed.

### Sucrose gradients

Continuous sucrose gradients were manually prepared. 50% and 15% sucrose solutions were prepared in RNAse free buffer (10 mM HEPES-KOH pH7.5, 150 mM KCl, 10 mM MgCl_2_, 100ug/mL cycloheximide and 1 mM DTT in RNAse free H_2_O). 1 mL each of decreasing concentrations of sucrose buffers, starting with 50% and topping off with 15% for a total of 11 mL was added to thin wall polypropylene ultracentrifuge tubes (Beckman Coulter). The tubes were immediately frozen and stored at -80 °C. On the day of use, the tubes were thawed at room temperature, resulting in the formation of a continuous gradient and immediately placed on ice. Formation of a continuous gradient was confirmed by adding different color dyes to the 50% and 15% starter sucrose solutions in a test preparation.

### Polysome fractionation

Polysome fractionation was performed as outlined previously (51). Briefly, roughly 80% confluent 10 mm plates of HEK293T cells was transfected with the *in vitro* transcribed test mRNAs. Two hours post transfection the cells were treated with media containing 100 ug/ml cycloheximide for 10 minutes to immobilize the ribosomes. The treated cells were then washed once with PBS containing 100 µg/ml cycloheximide. Cells were lysed on plate and harvested by adding 1 mL ice cold lysis buffer (10 mM HEPES-KOH pH7.4, 150 mM KCl, 10 mM MgCl_2_, 1mM DTT, 100 µg/mL cycloheximide, 2%NP-40, protease inhibitor, 6 U/mL RNase inhibitor). Lysates were centrifuged at 13000 rpm for 10 min at 4C to remove cellular debris. 800 µl of the lysate was loaded onto the sucrose gradient. The tubes were ultracentrifuged in a Beckman Coulter SW41 Ti swinging-bucket rotor at 40,000 rpm for 2 h at 4C. Samples were fractionated by pumping into the tubes a chase solution of 60% sucrose in RNAse free water using a BR-188 Density Gradient Fractionation System (Brandel). Real time concentration of RNA in the flow was measured by reading absorbance at 260 nm. 1 mL fractions were collected and set aside for RNA extraction. RNA was extracted from 200 µl of each fraction with Trizol reagent, and the quantity of transfected RNA in each fraction was determined by qRT-PCR. Absorbance data was analyzed by Peak Chart Data Acquisition software (Brandel) and overlaid with qRT-PCR data using Prism software (Graphpad).

## Acknowledgements

This work was supported by NIH grants AI120943 (Amarasinghe) and AI148663. We thank Benjamin tenOever (Icahn School of Medicine at Mount Sinai, New York, NY) for providing the RNAse III knockout cell line.

